# A literature review at genome scale: improving clinical variant assessment

**DOI:** 10.1101/193870

**Authors:** Christopher A. Cassa, Daniel M. Jordan, Ivan Adzhubei, Shamil Sunyaev

## Abstract

Introduction: Over 150,000 variants have been reported to cause Mendelian disease in the medical literature. It is still difficult to leverage this knowledge base in clinical practice as many reports lack strong statistical evidence or may include false associations. Clinical laboratories assess whether these variants (along with newly observed variants that are adjacent to these published ones) underlie clinical disorders.

Materials and Methods: We measured whether citation data—including journal impact factor and the number of cited variants (NCV) in each gene with published disease associations—can be used to improve variant assessment.

Results: Surprisingly, we find that impact factor is not predictive of pathogenicity, but the NCV score for each gene can provide statistical support of pathogenicity. When combining this gene-level citation metric with variant-level evolutionary conservation and structural features, classification accuracy reaches 89.5%. Further, variants identified in clinical exome sequencing cases have higher NCV scores than simulated rare variants from ExAC in matched genes and consequences (p<2.22x10^-16^).

Discussion: Aggregate citation data can complement existing variant-based predictive algorithms, and can boost their performance without accessing and reviewing large numbers of manuscripts. The NCV is a slow-growing metric of scientific knowledge about each gene’s association with disease.

Funding: This research was supported by NIH NHGRI grant HG007229 (C.C.) and NIGMS grant GM078598 (I.A., D.J., and S.S.).

## Introduction

Vast numbers of genetic variants have been reported to cause disease in the medical and scientific literature^1^, but it is still difficult to leverage this knowledge base clinically^2^. Variants that are identified in well-characterized genes such as *BRCA2* or *LDLR* would naturally warrant further review and potential clinical surveillance^3^. However, there are now approximately 5,000 genes with some clinical significance, many of which lack strong statistical evidence for pathogenicity, which we hereafter colloquially refer to as ‘disease-associated’. The majority of these genes have only a few publications describing their clinical significance^2^. Patients undergoing clinical genomic sequencing often carry novel variants in these less-characterized disease associated genes^4^, leading to challenges in diagnosis and complications in medical management^5^.

This dearth of information makes it difficult to translate the presence of a variant into an estimate of disease risk, particularly in carrier testing for asymptomatic individuals or in patients with multiple phenotypes where the range of associated disease genes is unclear^6–8^. When there are no reports describing an identified variant, clinicians must rely primarily on computational predictive techniques and published gene-level evidence to predict its functional effect^2^.

Even in more established disease genes, the majority of cases contain variants that are observed only in a single family, and labs must routinely prioritize and classify these variants of unknown significance^9,10^. Further, when specific variants have been previously described in publications, the knowledge base may be inadequate for clinical interpretation. Many of the variants that are in publication databases such as HGMD were originally identified in small, symptomatic populations without matched control groups, so their associations can suffer from incorrect estimates of significance or effect size, and a non-trivial fraction are likely to be spurious^11,12^. The result is a mixture of well-established associations with unverified or even false-positive findings.

For these reasons, a central challenge in clinical genomics is to prioritize variants that are identified during sequencing. We address this by integrating publication support for disease association at the gene level with variant-level estimates of functional impact to improve the variant assessment process.

## Citation-based predictive features

While estimates of effect size or significance may be missing or unreliable in case reports, it is difficult to ignore the clinical significance of many existing associations^13–15^. To address the uncertainty in individual reports, we extract information from publications—in aggregate across suspected disease genes—to assess the relative strength of each gene’s association with disease. The goal is to identify a metric that would have broad utility in separating highly suspicious genes from those which are more weakly associated with disease in large gene panels and clinical exome interpretation.

We evaluate two citation-based features in variant-level assessment:

1. **Number of cited variants (NCV) score**. Using a large set of published disease associations (HGMD 2012.1)^1^, we calculate the number of distinct variant sites in each gene that are described as causal.
2. **Impact factor (IF) score**. Each variant in HGMD is associated with a single publication, from which we identify the journal impact factor (Thomson Reuters)^16^. The impact factor is defined as the average number of citations over the prior two years for all manuscripts in a journal^17^.

To evaluate whether these features are predictive of pathogenicity, we apply each score to a large set of putative disease-associated variants. For each variant in HGMD 2012.1 (restricted to “DM” variant class; importantly, this version was not explicitly filtered by variant frequency), we calculate the allele frequency from a large population cohort (Exome Aggregation Consortium, ExAC, N=60,706 individuals). We then compare the NCV and IF scores for each variant with its observed allele frequency. In this set of disease-associated variants, we expect that variants with higher population frequencies are less likely to be highly penetrant and pathogenic, and those with lower population frequencies are more likely to have stronger association with disease^18^.

Surprisingly, journal impact factor is uncorrelated with variant allele frequency, indicating that variants reported in high-impact journals, on average, are not more likely to be pathogenic (**Figure 1A**, Supplementary Materials). However, we find a significant inverse correlation between NCV and variant allele frequency in putative disease variants. This suggests that variants in genes with lower NCV scores are less likely to be highly-penetrant pathogenic variants (**Figure 1A**). This trend is observed when restricting to rare variants (AF < 0.01) as well as a broader set of variants (AF < 0.05, Supplementary Materials).

**Figure 1:**
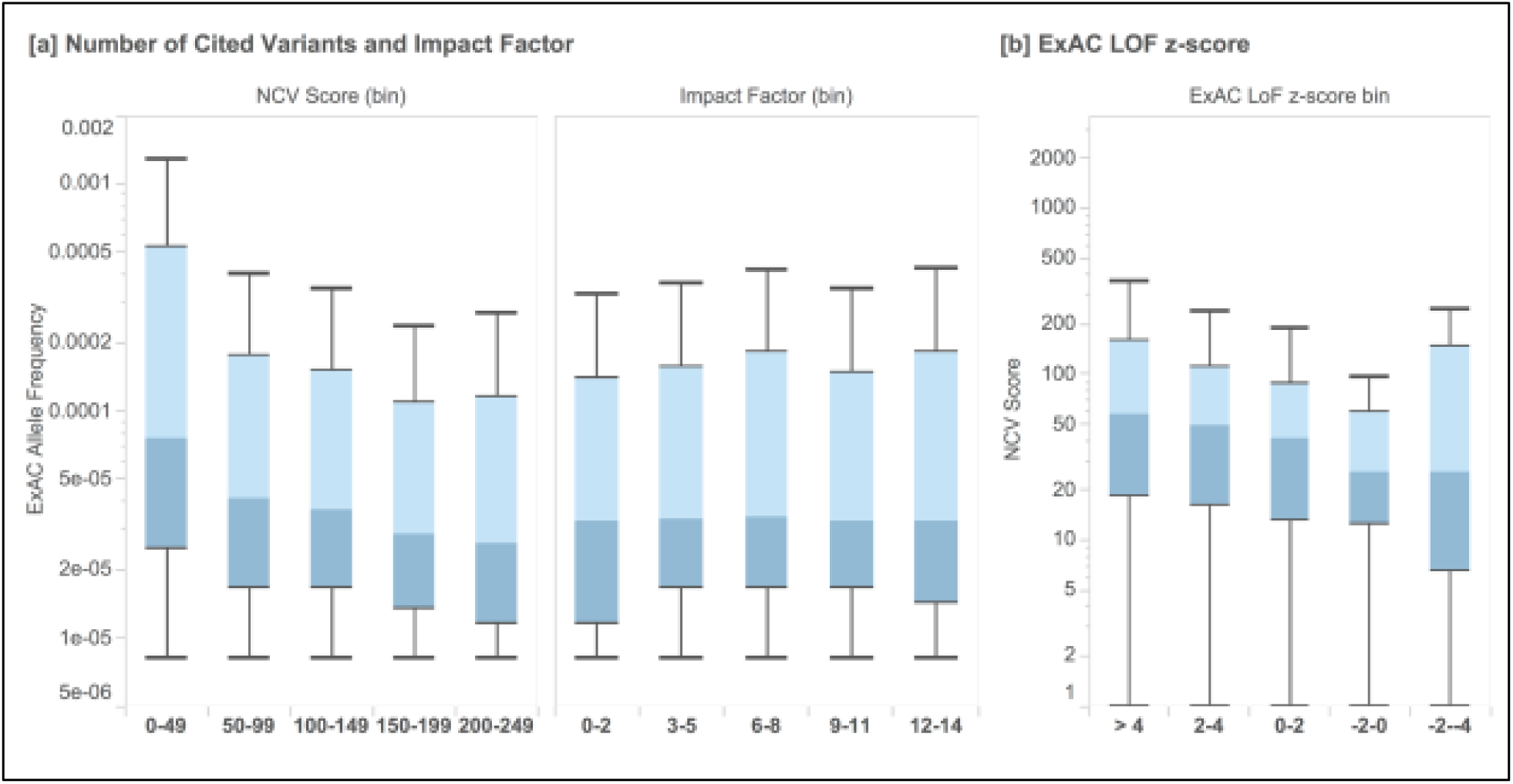
**[a]** Box plots of the allele frequency (Exome Aggregation Consortium, N=60,706) of a set of putative disease variants (HGMD 2012.1), grouped by the number of cited variants (NCV) in each gene and the journal impact factor for each citation. We observe an inverse correlation between the NCV score and allele frequency, while the impact factor of the journal in which a variant is cited is uncorrelated with allele frequency. **[b]** We find that genes under stronger selective constraint (those with positive z-scores reflect genes that have fewer loss of function [LoF] variants than expected based on a mutational model) have higher NCV scores than those genes which are already associated with disease but under lower selective constraint (p-value = 2.36x10^-3^).

At the gene level, we also find that the NCV score correlates well with an independently ascertained measure of genic intolerance to variation^19^. Disease-associated genes that are under stronger selective constraint (as estimated by the depletion of loss of function variants from expectation) have higher NCV scores than disease-associated genes that are under neutral or relaxed constraint (**Figure 1B,** Mann-Whitney U p-value = 2.36x10^-3^).

## Predicting variant pathogenicity in clinical exome sequencing cases

Next, we measure the performance of the NCV score in discriminating variants described as causal in clinical exome sequencing cases (Baylor Genetics)^9^. We compare the distribution of observed NCV values in Baylor cases with a set of realistic simulated “candidate” variants. The variants are drawn randomly from the same set of disease genes based on their ExAC allele frequencies^19^, restricted to variants below 0.1% and specific functional effects (**Supplementary Materials**). We find that causal Mendelian variants from this clinical exome sequencing program have significantly higher NCV scores than simulated variants (**Figure 2A**, p<2.22x10^-16^). This demonstrates that NCV can significantly improve standard filtering and prioritization by allele frequency in clinical exome sequencing cases.

**Figure 2:**
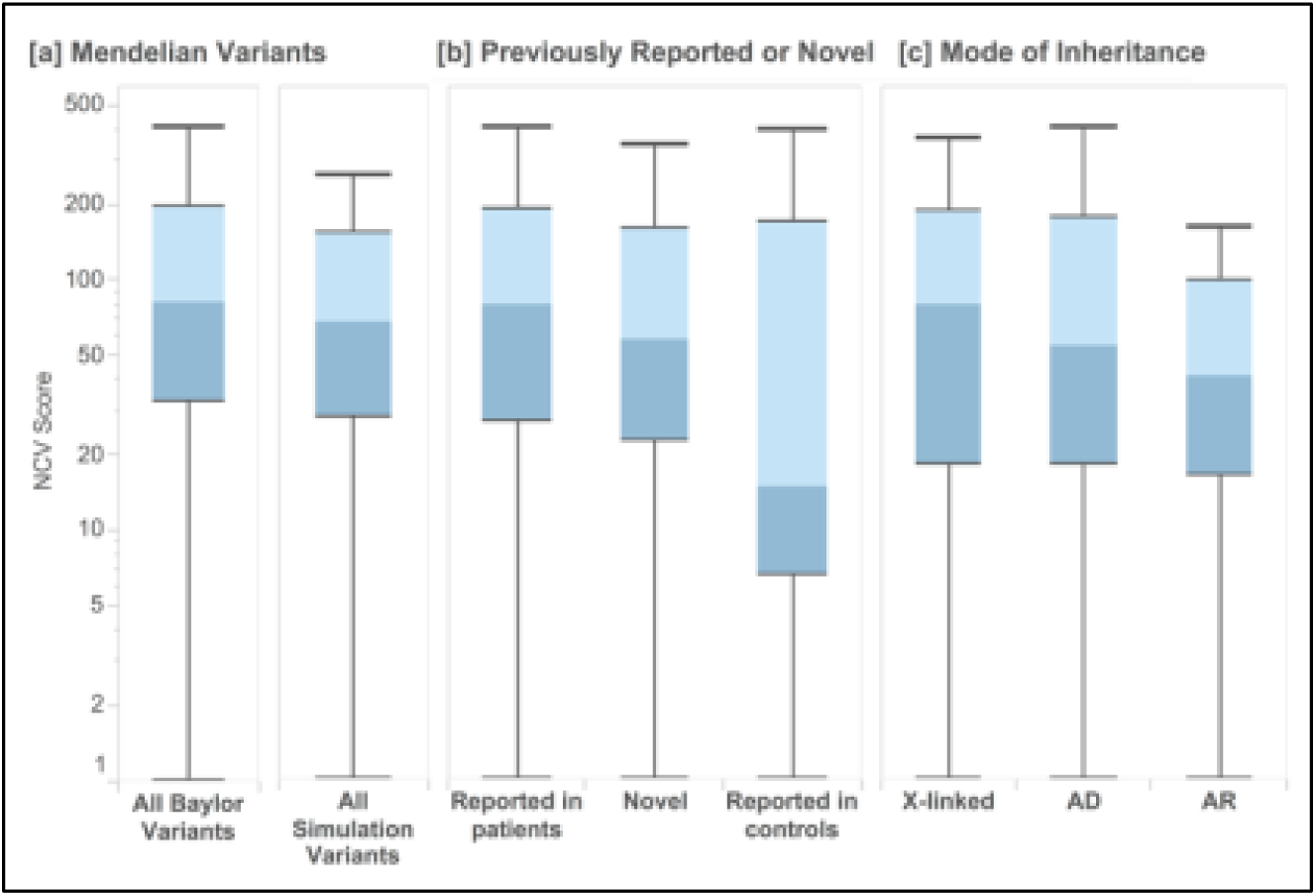
NCV score applied to clinical exome sequencing cases from Baylor Genetics and from simulated individuals. **[a]** Variants ascertained from clinical exome sequencing cases which resulted in a molecular diagnosis of Mendelian disorder had significantly higher NCV scores than those simulated from individuals in ExAC (Mann-Whitney U p-value < 2.22 x 10^-1^^6^). **[b]** Variants identified as causal in clinical sequence interpretation which had been previously reported in patients had higher NCV scores, followed by novel variants in known genes, and by variants which are also seen in control populations. **[c]** By mode of inheritance, cases with X-linked inheritance had higher NCV scores than those associated with autosomal dominant and autosomal recessive inheritance.

As expected, variants identified in clinical sequencing which had previously been reported as causal in clinic patients had significantly higher NCV scores than those that were previously reported in control populations (p=1.26x10^-16^) (**Figure 2B**). We also find that X-linked recessive variants from Mendelian cases have higher NCV scores than autosomal dominant (p=3.51x10^-4^) and autosomal recessive cases (p=7.77x10^-4^, **Figure 2C**). This may be due to differences in the ability to identify recessive disorders, differences in the genetic heterogeneity of recessive disorders, or another unknown source of bias.

## Integrating the NCV score with variant-level predictions of functional impact

Next, we sought to determine whether the gene-level NCV score provides complementary information to predictions made using structural and evolutionary features at the variant level. We first use the NCV score to classify the pathogenicity of individual variants, a common requirement in clinical interpretation. We apply the NCV score to each variant in HumVar, a gold-standard test dataset that contains missense variants which are already classified as either benign or pathogenic, derived from UniProtKB-Swiss-Prot,^20^ often used in classifier training^20^. We make extensive efforts to prevent bias and overfitting in our predictions^21^ (**Supplementary Materials and Supplementary Table 1**).

We find that the NCV score in each gene has predictive value, and can classify HumVar variant pathogenicity with an AUC of 0.847 (Naïve Bayes classifier in 5-fold cross-validation) (**Figure 3, green**). These gene-level predictions of pathogenicity are provided alongside NCV values in Supplementary Table 2, and an online interface [<u>http://genetics.bwh.harvard.edu/genescores/</u>].

**Figure 3:**
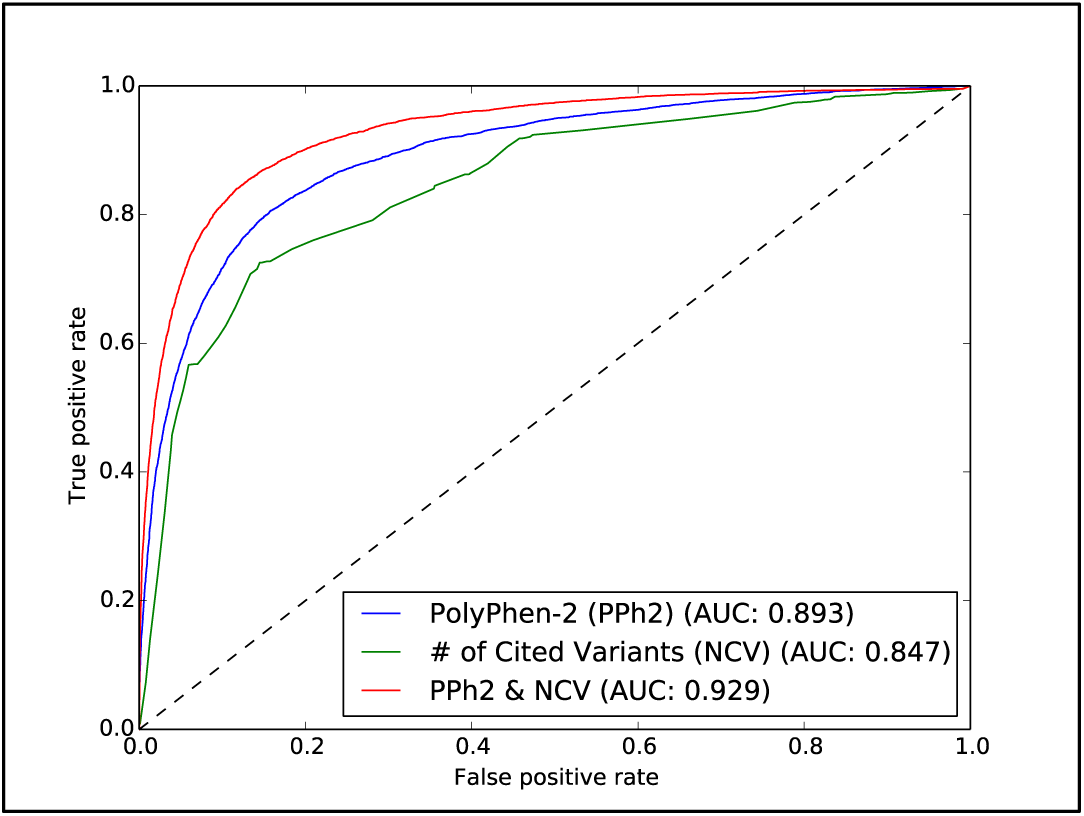
Receiver Operating Characteristic (ROC) curves from Naïve Bayes classifiers trained on the same gold-standard dataset from UniProtKB (HumVar). The number of cited variants (NCV) score is generated using a fully disparate set of citation data from HGMD variants that are not recorded in UniProtKB. A classifier trained using PolyPhen-2 features (blue) is compared with one that uses only the NCV score (green). Combining the NCV score with PolyPhen-2 features improves prediction accuracy substantially (red).

We then supplement this gene-level citation feature with individual variant-level predictions made by PolyPhen-2, a variant classifier that uses evolutionary conservation and structural features. We integrate these scores together using the probability of pathogenicity output by PolyPhen-2 as a prior for our own Naive Bayes NCV gene score, producing a composite score (**Supplementary Table 1**). The existing performance of PolyPhen-2 on the same dataset (without NCV) has an AUC of 0.893 (**Figure 3, blue**). When combining the gene-level NCV score with PolyPhen-2 features, the AUC on the same dataset rises to 0.929 (**Figure 3, red**), a substantial improvement in classification performance.

We find that the accuracy of this approach is not generated by a small set of well-known disease genes. We remove all variants appearing in any of the 56 genes included in the ACMG guidelines^3^—accounting for 1,999 of the 28,016 variants (7.14%)—and observe only a small reduction in AUC in our test dataset (0.8586 to 0.8575). We also find only a small effect when removing genes with outlier NCV scores (Supplementary Materials).

## Lower NCV scores observed for dosage sensitive, severe disease genes

Interestingly, we find the opposite effect in genes under the very strongest selection, where even the loss of one copy results in a severe phenotype. In a set of high-confidence haploinsufficient autosomal disease genes (N=127)^22^, we find that that genes with the strongest phenotypic severity, earliest age of onset, and highest fraction of *de novo* variants are associated with lower NCV scores (**Figure 4A-C**). This is consistent with previous work which demonstrated that genes under extremely strong selection (*e.g.* genes with increased embryonic lethality in mouse, cell essentiality, and severely depleted of variation in human populations) have fewer publications in PubMed than genes which are associated with weaker selective effects^23^.

**Figure 4:**
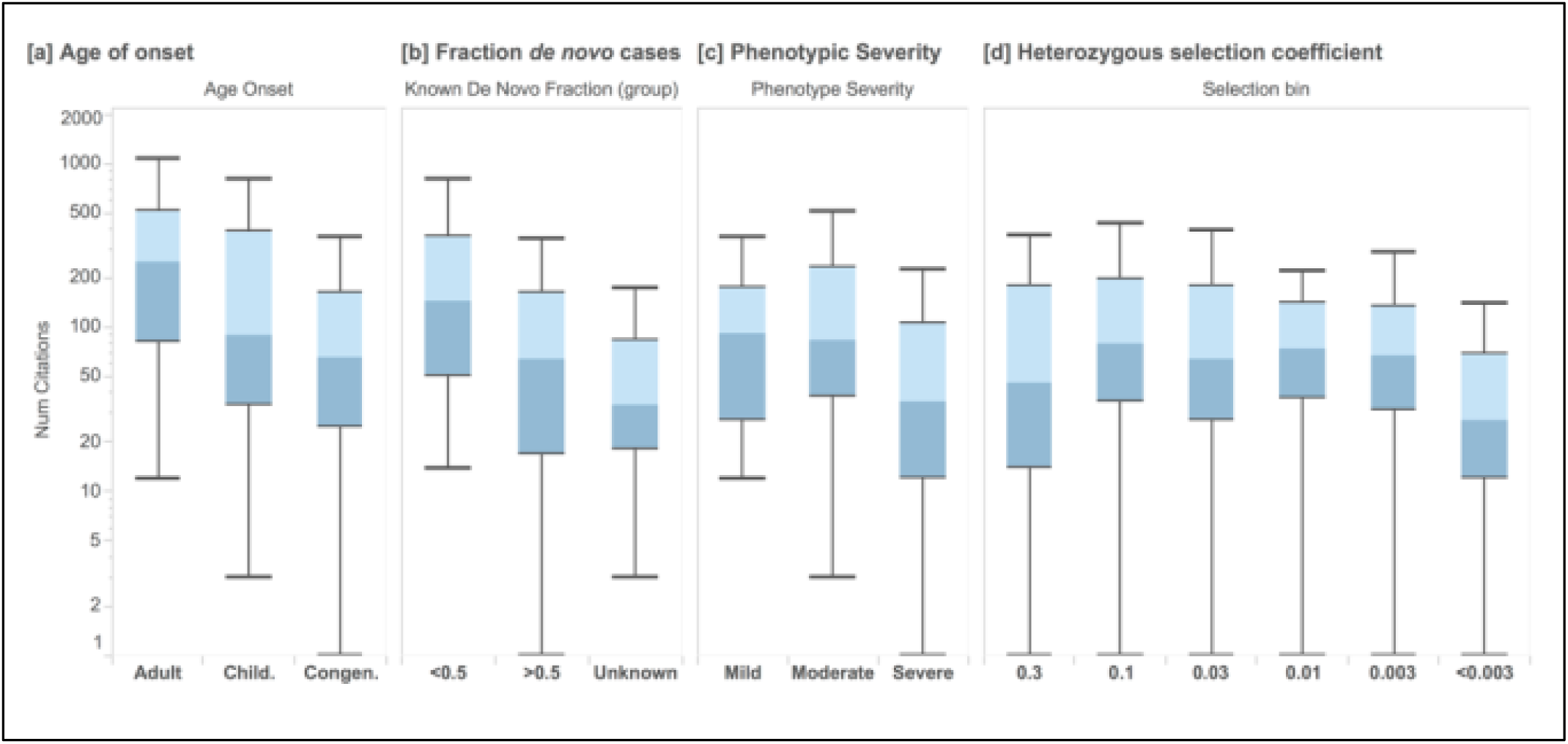
NCV score applied to haploinsufficient disease genes (ClinGen Dosage Sensitivity Project). In (N=127) autosomal genes, we annotate each disease associated gene with its NCV score, for each disease category and classification. Higher NCV scores are associated with **[a]** earlier age of onset (Mann-Whitney U p=0.0139), **[b]** a larger fraction of de novo variants (p=0.00345), **[c]** increased phenotypic severity (p=0.00494). Box plots range from 25th-75th percentile values and whiskers include 1.5 times the interquartile range. **[d]** We find that genes similarly enriched with more severe clinical annotations.

## Discussion

In this study, we demonstrate that aggregate citation data can be used at both the gene and variant level to improve pathogenicity prediction of novel variants. We show that these data can be combined with existing variant-level classifications, and have potential clinical utility in classifying variants in rare disease cases along with other indicators of pathogenicity. This helps address a substantial issue in clinical sequence interpretation, where there are often several variants under consideration in each case.

Ultimately, when a variant of unknown significance is encountered, the steps to resolve which variant is causal are costly, and include an in-depth literature review of each gene, familial segregation and functional validation studies^6^. The NCV score can be used along with variant-level scores to prioritize which variants should be considered first for these deeper studies. While clinical labs have developed efficient variant assessment processes, it difficult to assess the accuracy or clinical relevance of all existing publications within a gene, or to understand the relative importance of published literature in one gene relative to all other genes. For this reason, it is useful to employ automated approaches to assist with prioritization, so that resource-intensive processes can be restricted at first to a small set of genes.

If a gene has many variant sites associated with disease and few benign sites, it is more likely that subsequently identified variants in that gene are associated with disease. The NCV score is a gene-based feature that captures the number of sites in a gene that are reported to be disease-associated versus neutral. We gain predictive value using the genic distribution both neutral and pathogenic variants in each gene in HumVar, and use these data to provide a posterior probability based on the NCV score for each gene^24,25^. While the NCV is a gene-level score, a high NCV score only increases our confidence that the gene itself has a well-documented role in disease, but many variants within well-known disease genes may not be causal. This is precisely where information from functional predictions at the variant level provide complementary information to the NCV score and can provide utility in variant assessment.

For example, if a variant is observed in *BRCA2* with a possibly-damaging PolyPhen-2 score, it would likely rise to a probably-damaging score level, because of the preponderance of variants in *BRCA2* that are pathogenic (66.8%) and the lack of neutral or benign variants in the gene. In this case, a variant identified in *BRCA2* would have its PolyPhen-2 variant-level score multiplied by 1.335 (scores in Supplementary Table 2), which would alter its prediction of pathogenicity. For other genes, the score change may be smaller, especially if the knowledge base for the gene is less comprehensive.

In this case, the NCV score provides an assessment of gene morbidity and the variant classification provides information about the impact of specific variants, differentiating between those that are likely to disrupt a specific functional region of consequence or disrupt protein stability and those that have little or no effect on protein function^26^. This is consistent with previous results showing that different genes have measurably different thresholds of pathogenicity, and the accuracy of prediction methods can be improved by applying different thresholds or priors to different genes^27^.

Interestingly, we find that impact factor, on average, is not predictive of pathogenicity, and serves as a negative control in this study (**Figure 1**). It may be the case that journal impact factor is simply too statistically noisy to predict pathogenicity, or that individual article citation data would correlate more closely than the journal impact factor. The NCV score is less susceptible to noise, as it is a slow-growing metric that reflects a broad perspective about the knowledge base for each gene, and outperforms other citation-based features.

There are many factors which may potentially influence the NCV score that are difficult to model without extensive data. Genetic tests that are commonly indicated (*e.g.* cancer, hearing loss, cardiomyopathy) based on disease prevalence will lead to increased testing and observation of variants in known genes, potentially overweighting the NCV score. Conversely, genes associated with autosomal recessive disorders, or disorders largely driven by gain of function mutations will both have lower NCV scores.

There are several potential limitations in this study. First, we did not attempt to remove review articles or identify articles that refute previous claims due to review complexity. We also recognize the feedback potential for variants that are classified as pathogenic merely because previous variants in that gene have been classified as such. These issues remain difficult to resolve in computational predictions^21^. While longer genes have an increased target size, we have made our comparisons with features that are unrelated to gene length (*e.g.* allele frequency, loss of function depletion, and clinical categories.) We specifically chose to use the number of cited variants rather than a proportion of variant sites normalized by gene length due to concerns related to genes that have large regions which are tolerant to variation yet only a small gain-of-function region associated with disease, but also provide length-normalized results (Supplementary Materials).

While many HGMD variants may not be high-penetrance Mendelian variants^28,29^, the literature contains many high-quality disease–gene associations^15,30^. This approach extracts aggregate value from the large number of previously reported disease associations whose uncertain significance would make clinical classification difficult.

## Acknowledgements

This research was supported by NHGRI grant HG007229 (C.C.) and NIGMS grant GM078598 (I.A., D.J., and S.S.). We thank HGMD for their permission to publish these aggregated data, and Dr. Anne O’Donnell-Luria for access to curated data related to disorders in the ClinGen Dosage Sensitivity Project.

